# Modeling vascular dynamics at the initial stage of endochondral ossification on a microfluidic chip using a human embryonic-stem-cell-derived organoid

**DOI:** 10.1101/2024.08.18.608259

**Authors:** Abhiraj Kesharwani, Shoichiro Tani, Masaki Nishikawa, Yasuyuki Sakai, Hiroyuki Okada, Shinsuke Ohba, Ung-il Chung, Hironori Hojo

**Author notes:** Corresponding author (H.H.).

## Abstract

Vascular interactions play a crucial role in embryogenesis, including skeletal development. During endochondral ossification, vascular networks are formed as mesenchymal cells condense and later invade skeletal elements to form the bone marrow. We and other groups developed a model of endochondral ossification by implanting human embryonic stem cell (hESC)–derived sclerotome into immunodeficient mice. However, *in vitro* models of endochondral ossification, particularly vascular interaction with mesenchymal cells at its initial stage, are yet to be established. Therefore, we developed a method to model the initial stage of endochondral ossification using a microfluidic chip-based platform, with a particular focus on the vascular interaction. On the chip, we found that the fibrin gel helped align mCherry-expressing human umbilical vein endothelial cells (HUVECs) better than the collagen-I gel, suggesting that the fibrin gel is more suitable for the formation of a vascular-like network. The perfusability of the vascular-like networks was partially confirmed using fluorescein isothiocyanate (FITC)-dextran and fluorescent microbeads. We then mixed hESC-derived sclerotome with enhanced green fluorescent protein (EGFP)-expressing HUVECs and applied this mixture on the chip. We named this mixture of cells SH organoids. The SH organoids showed superior abilities to maintain the vascular-like network, which was formed by the mCherry-expressing HUVECs, compared with the sclerotome spheroids on the chip. The EGFP-expressing HUVECs migrated from the SH organoid, formed a vascular-like networks, and partially interacted with the mCherry-expressing vascular-like networks on the chip. Histological analysis showed that SRY-box transcription factor 9 (SOX9) and type I collagen were expressed mutually exclusively in the condensed mesenchymal cells and perichondrial-like cells, respectively. This study demonstrates that our SH organoid-on-a-chip method reproduces vascular networks that are formed at the initial stage of endochondral ossification. This model may provide insights into human endochondral ossification and has potential applications in bone disease modeling and drug screening.

## 1. Introduction

The vascular system is among the first organ systems to function during embryonic development. It forms a highly organized and continuous network that is crucial for the delivery of oxygen and nutrients to developing organs, playing vital roles in organogenesis. Endothelial cells, which line blood vessels, form the foundation of the vasculature. Far more than passive channels for blood flow, endothelial cells actively regulate tissue-specific development, shape organ formation, and influence overall organ function, possibly through paracrine or juxtacrine signaling mechanisms. [1–3].

Vascular formation is tightly coordinated with skeletal development. The vascular network appears as mesenchymal cells condense during the initial stages of skeletal development. In endochondral ossification, one of the two ossification modes, condensed mesenchymal cells differentiate into SOX9-positive chondrocytes and type I collagen-positive perichondrial cells, which are fibroblastic cells within the thin-layered perichondrium. Subsequently, chondrocytes form a cartilage mold, around which vascular networks form. Later, chondrocytes differentiate into hypertrophic chondrocytes, which serve as scaffolds for vascular invasion, initiating bone marrow formation[4–5].

Because mechanisms underlying interactions between vascular networks and skeletal development in humans have not been well understood, recapitulating the interaction using human embryonic stem cell (hESC) is highly desired [6–8]. We recently studied gene regulatory mechanisms in skeletal development using a hESC model, in which hESC-derived sclerotome cells were implanted underneath the renal capsule of immunodeficient mice [9]. We observed that the implanted sclerotome self-assembled and interacted with blood vessels to form the bone tissue. However, this system requires an *in vivo* mouse environment, which allows mouse cells to construct blood vessels in the hESC-derived bone; the interaction between human and mouse cells does not necessarily recapitulate corresponding events in human development. Therefore, we need an *in vitro* fully human model that recapitulates vascular interactions with skeletal cells during initial stage of endochondral ossification by using only human cells.

Recently developed organoid method and organ-on-a-chip technology have enabled more precise recapitulation of *in vivo* development. An organoid is a self-organized three-dimensional (3D) tissue consisting of multiple cell types derived from hESCs or other cells, mimicking biological functions of an organ [10–11]. This technology provides a valuable platform for modeling human organ development and investigating various diseases [12]. An organ-on-a-chip is a system engineered using microfluidic technology, where microfluidic chips are designed to recapitulate the 3D microstructures of tissues [13–15]. These technologies have been used to model several organs including vascular networks. However, modeling vascular interactions with skeletal cells at the initial stages of endochondral ossification has not yet been reported. Since sequential skeletal development involves multiple cell types, recapitulating this process in an *in vitro* system remains challenging.

In this study, we aimed to model the vascular interactions with skeletal cells using a hESC-derived sclerotome and an organ-on-a-chip method. We optimized a culture condition on a microfluidic-based chip and developed a method to form vascular networks between blood vessel-like tissues along with endochondral ossification induced by the hESC-derived sclerotome.

## 2. Methods

### 2.1. Human embryonic stem cell culture and maintenance

We conducted experiments involving hESCs, adhering to the Ministry of Health guidelines and obtaining Ethics Committee approval from the University of Tokyo (#19-203). As previously described, we used the SEES3 hESC line [16]. The initial adaptation and maintenance of hESCs were carried out in a commercially available xeno-free culture system, utilizing StemFit AK02N medium (#AK02N, Ajinomoto), and dishes were coated with 5 ug/mL recombinant human vitronectin (#A14700, Gibco). The cells were cultured under optimal conditions at 37 °C with 5% CO_2_ in a humid environment. For efficient cell dissociation, 0.5 M ethylenediaminetetraacetic acid (EDTA) (#15575-020, Gibco) was diluted to 0.5 mM using Dulbecco’s Phosphate-Buffered Saline (DPBS) (#049-29793, Wako). To ensure enhanced cell viability during seeding or passaging, 10 μM Y-27632 (#034-24024, Fujifilm Wako) was used to prevent apoptosis.

### 2.2. Sclerotome differentiation of hESC under xenofree conditions

Subconfluent, undifferentiated hESCs were gently dissociated into small clumps by approximately 10 rounds of pipetting. These cells were then sparsely passaged at a ratio of 1:16 to 1:24 depending on the size of the hESC colonies. The passaging was performed onto new vitronectin-coated culture plates using StemFit AK02N medium supplemented with 10 μM Y-27632 overnight. It is crucial to sparsely seed the hESCs before differentiation to minimize the risk of cellular overgrowth, particularly during prolonged differentiation periods. Differentiation into sclerotomes was systematically conducted in a serum-free, xeno-free, feeder-free monolayer environment using a chemically defined B27/ITS medium (BIM) [9]. The BIM was prepared and sclerotome induction was performed following a previously established methodology [9]. In brief, BIM was prepared by combining 1% B27 Supplement Xeno-Free (#A1486701, Thermo Fisher Scientific), 1% ITS Liquid Media Supplement (#I3146, Sigma-Aldrich), 1% MEM Non-Essential Amino Acids Solution (#11140050, Thermo Fisher Scientific), and 55 μM 2-Mercaptoethanol (0.1% of 55 mM 2-Mercaptoethanol in DPBS) (#21985023, Thermo Fisher Scientific) in Dulbecco’s Modified Eagle Medium/Nutrient Mixture F-12 (DMEM/F12) (#11330032, Thermo Fisher Scientific). The hESCs were differentiated into sclerotomes (Scls) using a stepwise protocol (Fig. S1a): induction from hESCs to primitive streak by treatment with 5 μM CHIR99021 (#4423, Tocris); induction from primitive streak to paraxial mesoderm by treatments with 5 μM CHIR99021, 1 μM A 83-01 (#2939, Tocris), and 0.25 μM LDN193189 (#SML0559; Sigma); induction from paraxial mesoderm to somitic mesoderm by treatments with 1 μM C59 (#C7641-2S; Cellagen Technology), 1 μM A 83-01, and 0.25 μM LDN193189; induction from somitic mesoderm to sclerotome by treatments with 1 μM SAG (#AG-CR1-3585, Adipogen), 1 μM C59, and 0.25 μM LDN193189. All inductions were performed for 24 h, except for sclerotome induction, which lasted for 48 h.

### 2.3. Construction of a lentivirus vector

The pLenti-mCherry-Puro vector was constructed using the HiFi assembly kit (#E2621S, NEB) with an mCherry-T2A-PuroR (puromycin resistant gene) fragment into a lentiviral vector. The backbone of the lentiviral vector was prepared by removing the mCherry fragment from the pLV-mCherry vector (Plasmid #36084, Addgene).

### 2.4. Lentivirus preparation

HEK293T cells (#CRL-3216, ATCC) were cultured in Dulbecco’s Modified Eagle’s medium (DMEM), supplemented with 10% fetal bovine serum and 1% penicillin/streptomycin (#P4333, Sigma-Aldrich). The maintenance of cells occurred in a tissue culture incubator at 37 °C with 5% CO2. Upon reaching 60-70% confluency, HEK293T cell transfection was carried out using FuGENE® HD (#E2312, Promega) and Opti-MEM (#31985-070, Thermofisher) following the manufacturer’s instructions. Plasmids for vascular endothelial growth factor A (VEGFA)-EGFP lentivirus production were pCCLc-MNDU3-VEGFA-PGK-EGFP-WPRE (Plasmid #89609, Addgene), pMD2.G (Plasmid #12259, Addgene), and psPAX2 (Plasmid #12260, Addgene). For mCherry lentiviral production, the pmCherry-puromycin vector, pMD2.G, and psPAX2 were used. FuGENE HD reagent and Opti-MEM were incubated for 30 min at room temperature. For a single 10 cm dish, a mixture was prepared comprising 8.5 μg of pCCLc-MNDU3-VEGFA-PGK-EGFP-WPRE for VEGFA-EGFP transfection, or alternatively, 8.5 μg of pmCherry-Puro for mCherry transfection, along with 2.1 μg of pMD2.G, and 6.4 μg of psPAX2. The volume was adjusted by adding Opti-MEM to reach 799 μL, and then 51 μL of FuGENE HD transfection reagent was introduced to achieve the final volume of 850 μL. After combining the plasmids and reagent at room temperature, the mixture was added to HEK293T cells in a 100 mm dish containing 10 mL DMEM with 10% fetal bovine serum and 1% penicillin and streptomycin (#168-23191, Fujifilm Wako Pure Chemical). Following 48 hours of transfection, the cell culture medium was harvested and filtered using a 0.45 μm filter. For lentiviral storage and concentration, a Lenti-X concentrator (#631231; Takara) was used according to the manufacturer’s protocol. A concentrated lentivirus suspension was used to transduce HUVECs.

### 2.5. HUVECs lentiviral transduction

HUVECs (#00191027 XL size, Lonza) were cultured in Endothelial Growth Medium-2 (EGM-2) (#CC-3162, Lonza) and maintained in a tissue culture incubator at 37 °C with 5% CO_2_. Lentiviral transduction was conducted the following day when HUVECs reached an optimal confluency range of 40–50%. A concentration of 6 μg/mL Polybrene (#107689-10G, Sigma) was included in the lentiviral transduction process. After 48–72 h of transduction, GFP expression from VEGFA-EGFP lentivirus transduction or RFP expression from mCherry lentivirus transduction was observed under a fluorescence microscope. Overexpression of VEGFA was validated using RT-qPCR.

### 2.6. SH organoid formation with SH (Sclerotome + HUVECs) method-

Sclerotome cells derived from hESCs and HUVECs overexpressing VEGFA were dissociated using Accutase (#AT104, Innova Cell Technologies) and a 2.5% trypsin-EDTA solution (#T4174-100mL, Wako), respectively. After counting, centrifuging, and collecting the cells, sclerotome cells and HUVECs were resuspended in Scl induction medium with 1 μM Y-27632 and EGM-2, respectively. For the formation of SH organoids, sclerotome cells and HUVECs were mixed in a 4:1 ratio in SH medium and 3 × 10^4^ cells per well were seeded into a 96-well ultra-low attachment plate (#7007, Corning). The SH medium was prepared by combining EGM-2 and the Scl induction medium with 1 μM Y-27632 in a 9:1 ratio. After the initial 24-hour incubation period, the culture medium was changed to the EGM-2 medium, and subsequently, the EGM-2 medium was changed every other day (Fig.

S1a). The sclerotome spheroids were generated by seeding 5 × 10^3^ cells per well in a 96-well ultra-low attachment plate with sclerotome induction medium containing 1 μM Y-27632. After a 24-hour incubation period, the culture medium was replaced with EGM-2 medium, which was subsequently refreshed every other day to maintain optimal conditions for spheroid development.

### 2.7. Mimetas OrganoPlate^®^ Graft plate

The OrganoPlate^®^ Graft plate (#6401-400-B, Mimetas), is a microfluidic platform comprising 64 chips, specifically designed to fit into a standard 384-well plate. Each chip contained three distinct channels: one center channel for gel loading and two peripheral channels for the culture medium. The OrganoPlate^®^ Graft was originally intended for vascularizing tumor spheroids [17], but here we chose SH organoids to utilize the available open access (with a 1 mm diameter) and to establish the vascular-like networks underneath the gel chamber. Interaction between the SH organoid and vascular-like networks was observed under a fluorescence microscope.

### 2.8. Gel loading, and vascular-like network formation-

For gel loading, fibrinogen (#F3879-100MG, Sigma) and thrombin (#T6884-100UN, Sigma) stock solutions were prepared at concentrations of 15 and 100 U/mL, respectively, in DPBS. Then fibrin solution was prepared by combining 300 μL of 15 mg/mL fibrinogen with 10 μL of 100 U/mL thrombin in DPBS, resulting in a final volume of 1000 μL. The final working concentrations of thrombin and fibrinogen in the fibrin solution were 1 U/mL, and 4.5 mg/mL respectively. Subsequently, 3 μL of the fibrin solution were immediately injected onto the inlet of the gel chamber following the addition of thrombin to the fibrinogen solution. It was visually confirmed that the fibrin solution completely filled the gel chambers. Next the OrganoPlate^®^ Graft was incubated at 37 °C for 15 min to facilitate fibrin gel crosslinking. To prevent the drying of the fibrin gel, 50 μL of Hanks’ balanced salt solution (HBSS) (#H8264-100ML, Sigma) was added to the gel inlet.

To initiate the formation of vascular-like networks, mCherry-expressing HUVECs were dissociated using a 2.5% trypsin-EDTA solution. After counting, centrifuging, and collecting the mCherry-expressing HUVECs, 2 μL of the cell suspension, with each microliter containing 10,000 cells, were added to each perfusion channel. Subsequently, 50 μL of EGM-2 was introduced into each perfusion inlet, and the plate was incubated at 37 °C with 5% CO_2_ for 2–4 h under stable conditions. After the attachment of cells in the perfusion channels, an additional 50 μL of EGM-2 was applied to each perfusion outlet to establish complete medium flow within the perfusion channels. Then the HBSS was aspirated from the gel inlet, and 50 μL of EGM-2 was introduced into the graft chamber. Following this, the plate was cultured for 2-3 days on a rocking stage (OrganoFlow, Mimetas, USA) with a 14° inclination and an 8-minute interval to allow bidirectional flow in the chip. These specific culture conditions were maintained to promote the formation of tube-like structures within the perfusion channels, in accordance with the recommendations of Mimetas for optimal vessel formation and maintenance. Subsequently, the device underwent stimulation with 50 μL of an angiogenic cocktail, comprising recombinant human endothelial growth factor (rhVEGF)-165 (#100-20, Peprotech) (37.5 ng/mL), recombinant human fibroblast growth factor-2 (rhFGF-2) (#100-18B, Peprotech) (37 ng/mL), recombinant human hepatocyte growth factor (rhHGF) (#11343413, ImmunoTools) (37.5 μg/mL), recombinant human monocyte chemoattractant protein-1 (rhMCP-1) (#11343384, ImmunoTools) (37.5 ng/mL), phorbol 12-myristate 13-acetate (PMA) (#P1585, Sigma) (37.5 ng/mL), and sphingosine 1-phosphate (S1P) (#S9666, Sigma) (250 nM), aiming to induce the formation of a vascular-like networks in the graft chamber. The angiogenic cocktail was changed every day, and the EGM-2 in the perfusion channels was changed every 2 days. The culture was continued on a rocker for 3-4 days for vascular formation.

### 2.9. SH organoid grafting in the chip

On day 14, an SH organoid within the size range of 500 to 700 μm was carefully chosen from the organoid culture maintained under EGM-2 conditions. Using a 200 μL-sized tip, the organoid was picked up and grafted into the chip’s graft chamber where a vascular bed was present. The positioning of the organoid was fine-tuned using a needle to ensure its placement in the central part of the chamber. Following organoid grafting, 50 μL of an angiogenic cocktail was introduced into the graft chamber, and this cocktail was replaced daily and EGM-2 in perfusion channels were changed on alternate days throughout the 7-day culture period in the chip.

### 2.10. Sprout perfusability

To assess the perfusability of the constructed vascular bed within the chip, we introduced a solution of fluorescent microbeads (#T7284, TetraSpeck™ Fluorescent Microspheres Sampler Kit, ThermoFisher) and 70 kDa FITC-dextran (#53471-1G, Sigma) on day 7 into both perfusion inlets following removal of the cultured EGM-2 medium. The fluorescent microbead solution comprised beads sized at 4 μm, each possessing four distinct excitation/emission peaks: 360/430 nm (blue), 505/515 nm (green), 560/580 nm (orange), and 660/680 nm (dark red), with a concentration ranging from 7×10 ^5 to 14×10 ^5 beads per milliliter in PBS. The green fluorescent microbeads were visualized for several minutes using a BZ-X700 fluorescence microscope (Keyence).

FITC-dextran solution was prepared by dissolving 0.5 mg 70 kDa FITC-dextran (0.5 mg) in 1 mL of the EGM-2 medium. This solution was added both peripheral channels of the chip and observations were performed for several minutes using a BZ-X700 fluorescence microscope.

### 2.11. Gel area and sprouting quantification

The gel area was quantified using Fiji (ImageJ), whereas the evaluation of sprouting involved tallying the number of vessels and measuring their lengths using the same software. The length of each vessel was determined by tracing from the edges of the perfusion channels (on both sides) to the endpoint within the extracellular matrix (ECM). To quantify the vascular area, only the vascular-like networks within the gel were analyzed and expressed as a percentage.

### 2.12. Histological analysis

Sclerotome spheroid and SH organoid samples were retrieved from the chip using a needle and fixed in 4% paraformaldehyde at 4°C for 1 h. Upon fixation, the samples underwent three washes with PBS at 4°C for 30 min each. Prior to the preparation of frozen blocks, the samples were immersed in 30% sucrose in PBS at 4°C overnight. Following this, the samples were embedded in O.C.T. compound (#4583, Sakura FinetekTM Japan) and frozen at -80°C. Frozen sections, 10 μm thick, were obtained using a CM3050IV cryotome (LEICA) and mounted onto slides (SCRE-01, Matsunami), followed by air-drying at room temperature before staining. Hematoxylin and Eosin (H&E) staining was performed for morphological analysis. The sections were immersed in hematoxylin for 4 min, rinsed in tap water for 10 min, and briefly in 95% ethanol. Subsequently, eosin staining was applied for 1 min. The sections were dehydrated with 100% ethanol and xylene for 5 min each, repeated twice and thrice, respectively, before mounting with Marinol (#20092, Muto). For immunohistochemistry, antigen retrieval was performed for matrix staining with 20 μg/mL Proteinase K (Recombinant) solution (#1567906, Nacalai) in PBS. Blocking was conducted using Blocking One Histo (#06349-64, Nacalai), then primary antibodies against CD31 (#11265-1-AP, Proteintech), SOX9 (#Ab5535, Millipore), and type I collagen (COL1) (#Ab6308, Abcam) were used, along with secondary antibodies: Alexa Fluor 647 anti-rabbit IgG (A21245, Invitrogen) and Alexa Fluor 546 anti-mouse IgG (#A11030, Invitrogen) for fluorescence visualization. All antibodies were used at a dilution ratio of 1:500. Bright-field and fluorescence images were captured using a BZ-X700 fluorescence microscope (Keyence) and a confocal microscope (LSM 880; Zeiss).

### 2.13. RNA extraction and reverse transcription-quantitative polymerase chain reaction (RT-qPCR)

Total RNA was extracted from each sample using ISOGEN (#311-02501, Nippon Gene) and the RNeasy Mini Kit (#74106, Qiagen). RNA quality and concentration were assessed using a NanoDrop ND-1000 spectrophotometer (Thermo Fisher Scientific). Then, 1 μg of total RNA was reverse-transcribed into complementary DNA (cDNA) using the ReverTra Ace™ qPCR RT Master Mix with gDNA Remover Kit (FSQ-301, Toyobo Co., Ltd.). For the PCR reactions, 2 μl of cDNA was used per reaction, with amplification carried out using the FastStart Universal SYBR Green Master Kit (04913 850001, Roche). RT-qPCR was performed on a 7500 Fast Real-Time PCR System (Applied Biosystems). GAPDH served as the endogenous human control, and relative mRNA expression levels were determined using the ΔΔCT method. The specific human primers used for gene expression analysis are as follows:

Human glyceraldehyde-3-phosphate dehydrogenase, GAPDH: forward 5’ GAAGGTGAAGGTCGGAGTCA 3’ reverse 5’ GAAGATGGTGATGGGATTTC 3’

Human Nanog homebox, NANOG: forward 5’ AACTGGCCGAAGAATAGCAA 3’ reverse 5’ TGCACCAGGTCTGAGTGTTC 3’

Human POU domain, class 5, transcription factor 1, POU5F1: forward 5’ GAAGGATGTGGTCCGAGTGT 3’ reverse 5’ GTGAAGTGAGGGCTCCCATA 3’

Human paired box gene 1, PAX1: forward 5’ CCGCAGTGAATGGGCTAGAGAA 3’ reverse 5’ TACACGCCGTGCTGGTTGGAG 3’

Human paired box gene 9, PAX9: forward 5’ AGCAGGAAGCCAAGTACGGTCA 3’ reverse 5’ CAGAAGGAGCAGCACTGTAGGT 3’

### 2.14. 3D imaging

Live 3D imaging of SH organoids cultured on a Mimetas OrganoGraft® chip was captured using a confocal microscope over time. To create the 3D images, z-stacks were generated and merged using ZEN 3.4 blue edition. Ortho images were created using the same software.

## 3. Results

### 3.1. Establishing a vascular-like network in a microfluidic chip

To mimic the formation and interactions of peripheral blood vessels, we aimed to establish vascular-like networks within a microfluidic chip system. We utilized the OrganoPlate^®^ Graft from Mimetas, a versatile multichip microfluidic platform containing 64 individual chips per plate (Fig. 1a). Each chip featured 2 main peripheral channels connecting wells A1 to B1 and A3 to B3 along with a gel chamber with a gel-loading inlet in well A2 and open access in well B2. The organoid can be positioned in the open-access gel in well B2 (Fig. 1a, 1b). The process of forming vascular-like networks is as follows: extracellular matrix (ECM) and endothelial cells are introduced through the gel inlet and both peripheral channels, respectively. After tube formation, angiogenic factors can be added to well B2 to induce sprouting from the peripheral channels. Following the establishment of a vascular-like network, the organoid can be introduced, allowing the observation of vascular-like networks with the organoid (Fig. 1b).

**Fig. 1.**
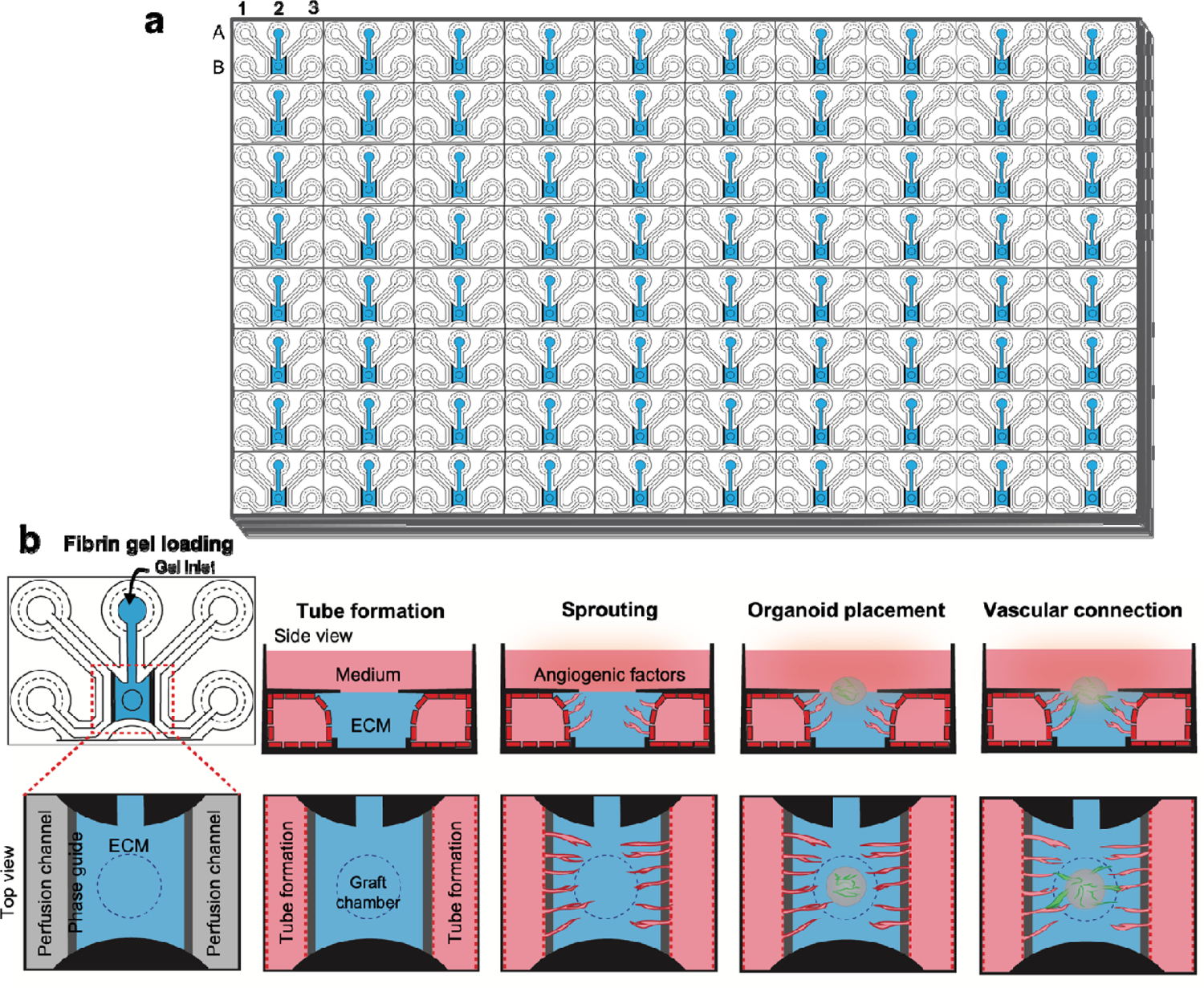
Schematic of organoid-on-a-chip in this study. (a) Illustration of the Mimetas OrganoPlate^®^ Graft, consisting of 384 wells and 64 chips. (b) Illustration of experimental steps for the blood vessels interaction into the organoid. Fibrin gel loading: the gel is loaded through the gel inlet. Tube formation: HUVECs are seeded in the perfusion inlets and vessel tube formation under flow. Sprouting: vascular-like network is formed using an angiogenic cocktail. Organoid placement: organoid is placed in a graft chamber. Vascular connections: vascular connection occurs between the organoid and vascular from peripheral channel.

To optimize the culture conditions for forming vascular-like networks on the chip, we evaluated 2 extracellular matrix (ECM) materials: collagen-I gel and fibrin gel. Both gels were prepared and loaded into the gel inlet of the chip. Following gel crosslinking, mCherry-expressing HUVECs were cultured in both perfusion channels of the chip for 2–3 days (Fig. 2a). Subsequently, an angiogenic cocktail containing VEGFA, FGF-2, MCP-1, HGF, S1P, and PMA were added to promote the formation of vascular-like network within the graft chamber (Fig. 2b). Morphological assessments showed that although both matrices exhibited minimal shrinkage prior to exposure to the angiogenic cocktail, the collagen-I gel shrank considerably compared with the fibrin gel (Fig. 2c). This caused detachment of the gel from the peripheral channels, resulting in no contact between peripheral channels and vascular-like networks. In contrast, the fibrin gel formed vascular-like networks in the presence of the angiogenic cocktail without significant shrinkage and kept the contact between peripheral channels and vascular-like networks. The number of vascular-like networks, and their lengths were significantly higher in the fibrin gel than in the collagen-I gel from days 4 to 7 (Fig. 2d). Additionally, vessel lengths were significantly increased in the fibrin gel compared to the collagen-I gel on days 6 and 7 (Fig. 2d).

**Fig. 2.**
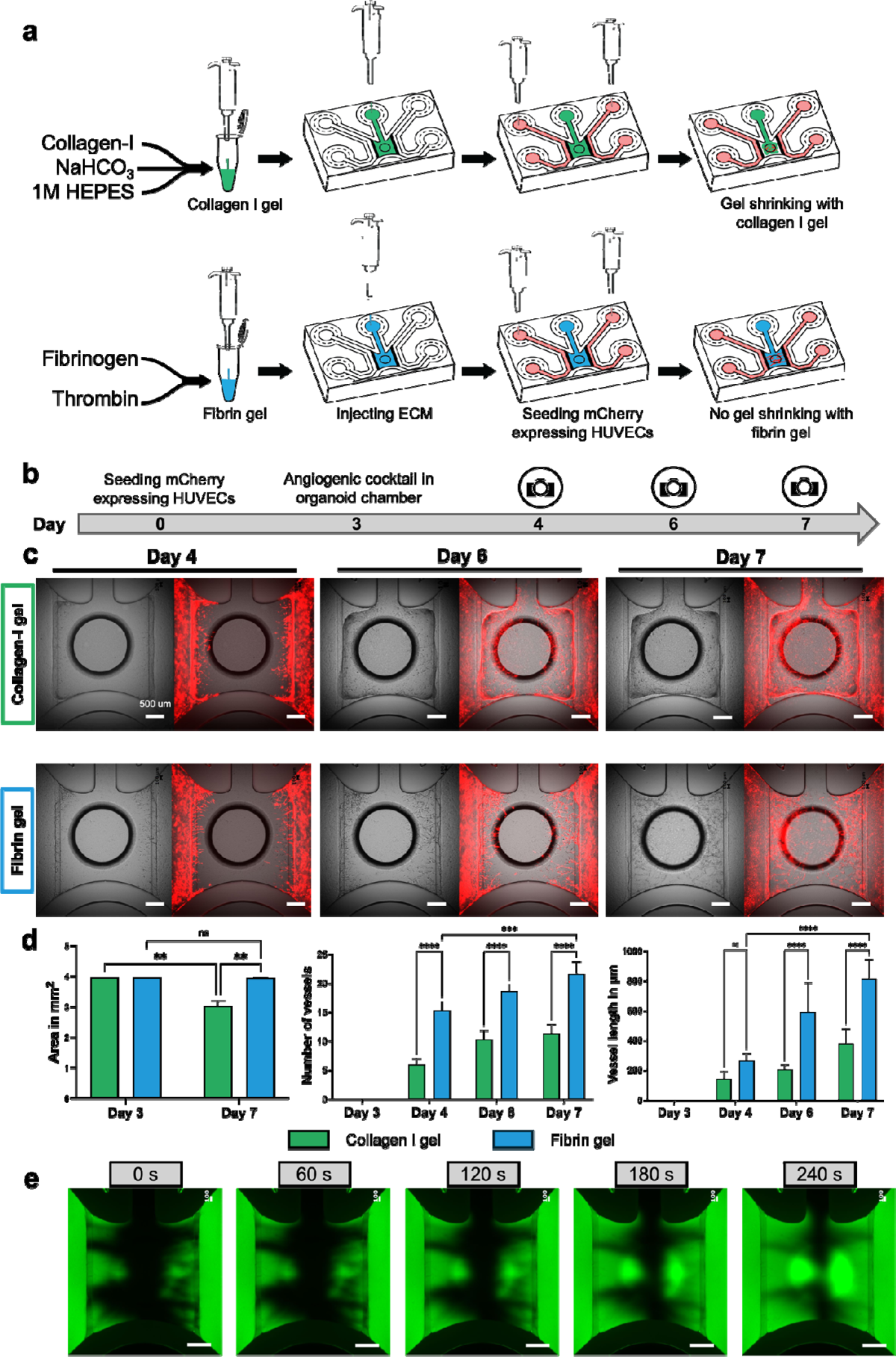
Development of perfusable vascular-like networks on the chip. (a) Representation of vascular bed formation with collagen-I and fibrin gel loading and HUVECs seeding in perfusing inlets. (b) Timeline for vascular bed formation. (c) Bright field and merged images represent the angiogenesis using collegen-I gel and fibrin gel. Scale bar: 500 μm. (d) Comparative analysis between collagen-I gel and fibrin gel in terms of gel area, number of vascular-like networks, and their lengths over 7 days of vascular-like network formation. A two-way ANOVA with multiple comparisons was used to analyze the data. Statistical significance is indicated by ** for p < 0.01, *** for p < 0.001, and **** for p < 0.0001. (e) Validation of vascular bed perfusibility in fibrin gel using 70 kDa FITC-dextran represented as a green fluorescent fluid for long-term observation. Scale bar: 500 μm.

Perfusability tests were conducted on the vascular bed within the fibrin gel. We introduced 70 kDa FITC-dextran into both perfusion channels. We found that perfusion began immediately after the introduction of FITC-dextran into the perfusion channels. At 0 s, FITC-dextran showed perfusion through the vascular bed and reached the center of the graft chamber during long-term observation until 240 s (Fig. 2e). Additionally, we found that the FITC-dextran solution did not perfuse or diffuse into the graft chamber in the absence of vascular-like networks on the chip (Fig. S2) For further perfusabiliy confirmation, a 4 μm size-fluorescent bead suspension was introduced into the perfusion inlets at 0 s. After 156 s, we observed fluorescent beads traversing the vascular-like networks within the chip. Hence, our findings indicate that fibrin gel exhibits superior qualities to collagen-I gel in the chip environment for aligning HUVECs, possibly providing better vascular-like networks on the chip.

### 3.2. Sclerotome spheroid and SH organoid on the perfusable vascular-like network and their interactions

To achieve vascular network interaction with skeletal cells in vitro, we hypothesized that mixing HUVECs with sclerotome that contains osteochondral progenitors would help form vascular-like networks on the chip. Using a previously published method [9], we differentiated hESCs into sclerotome and mixed them with HUVECs (Fig. S1a). Hereafter we called this mixture of cells the SH organoid (Sclerotome and HUVECs) and set day 0 as the day of the SH organoid formation (Fig. S1b). We confirmed sclerotome induction by RT-qPCR. It showed downregulated expression of the pluripotency-related genes *NANOG* and *POU5F1* and upregulated expression of the sclerotome-related genes *PAX1* and *PAX9* (Fig. S1b). To enhance the visibility of HUVECs, we labeled the cells with EGFP by lentiviral transduction and used the EGFP-expressing HUVECs for the SH organoid formation.

We compared the effect on vascular-like network formation between the SH organoid and sclerotome spheroids on the chip (Fig. S1b). Once the vascular-like networks fully developed within the graft chamber, the sclerotome spheroids or SH organoids were introduced into the graft chamber of the chip. We used the angiogenic cocktail for one week in the culture of the sclerotome spheroid and SH organoid to maintain vascular-like networks on the chip. During culture, we observed connections between mCherry-expressing HUVECs from peripheral channels and EGFP-expressing HUVECs from the SH organoid, suggesting the formation of vascular-like networks. (Fig. 3a). By day 14, the degree of the vascular-like networks were compromised to some extent under both conditions. To assess vascular conditions, we examined mCherry-expressing vascular-like network areas on days 8 (Fig. 3b, c) and 14 (Fig. 3d, e). Microscopic images and quantitative analysis showed comparable vascular-like networks under both conditions on day 8 (Fig. 3b, c). On day 14, although we observed degradation of the vascular-like networks under both conditions, the degree of the vascular-like networks was better in the SH organoid than in the sclerotome spheroid on the chip (Fig. 3d, e). These results indicate that the SH organoid has better abilities to maintain vascular-like networks than the sclerotome spheroids on the chip until day 14 of the culture period.

**Fig. 3.**
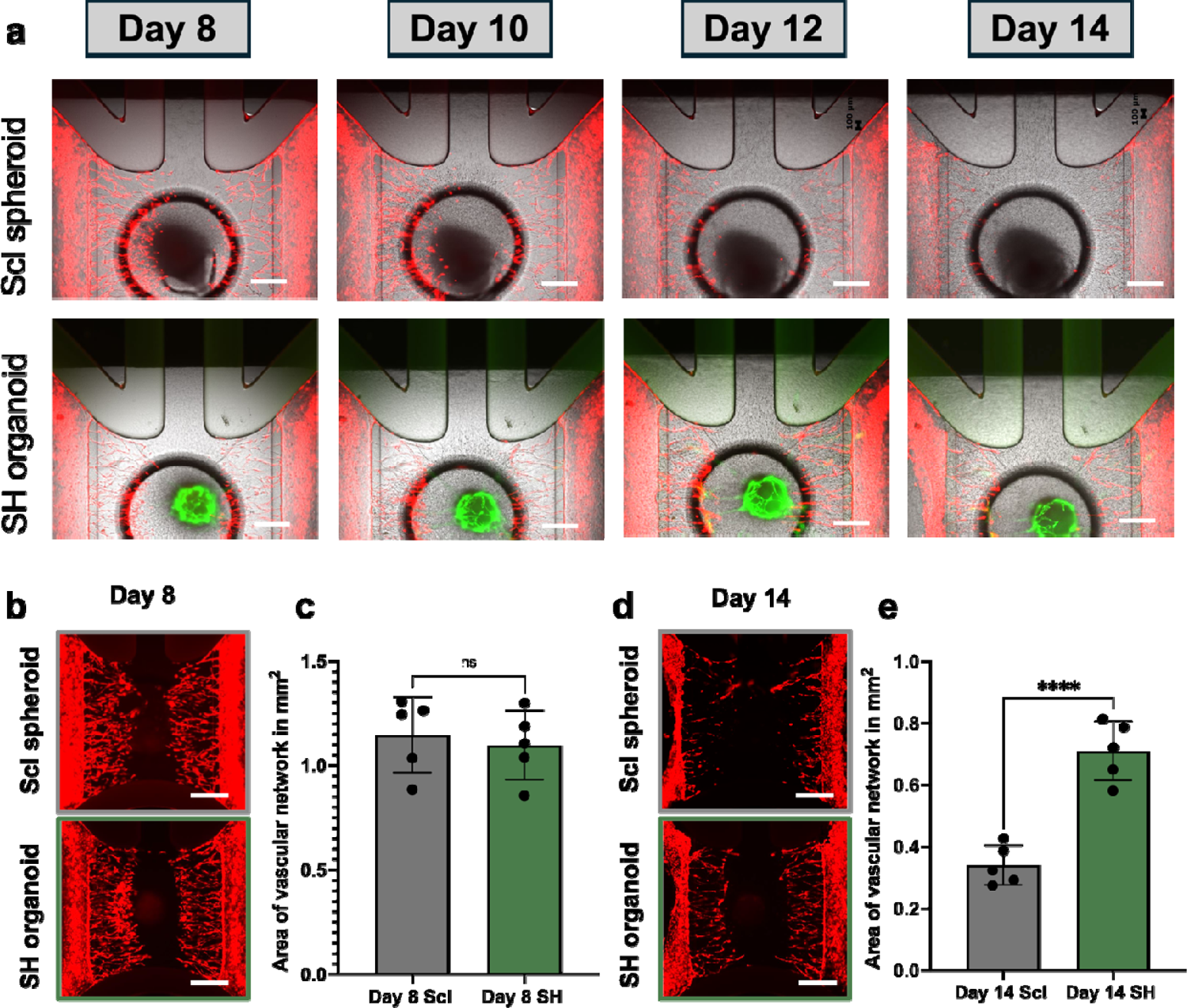
Sclerotome spheroid and SH organoid culture on a vascular-like network. (a) Fluorescent microscopic images of sclerotome spheroid- and SH organoid-on-a-chip captured from day 8 to day 14. (b, c) Fluorescent image of mCherry expressing vascular-like networks (b) and the quantification analysis (c) on day 8 for sclerotome spheroid- and SH organoid-on-a-chip. (d, e) Fluorescent image of mCherry expressing vascular-like networks (d) and the quantification analysis (e) on day 14 for sclerotome spheroid- and SH organoid-on-a-chip. In A, B, D, Scale bar: 500 μm. In c, e, Scl, sclerotome spheroid; SH, SH organoid. For statistical analysis Student’s t-test was used, “*” indicates statistical significance (****p < 0.0001) and “ns” represents not significant.

To assess the interaction between the SH organoid and the peripheral vascular-like network, we observed fluorescent microscopic images of the SH organoid-on-a-chip. We found that the SH organoids formed EGFP-expressing vascular-like networks within the matrix on day 8 on the chip, characterized by the emergence of small GFP-expressing vascular-like sprouts extending from the SH organoids (Fig. 4a). The vascular-like sprouts extended and reached the peripheral mCherry-expressing vascular-like networks by day 10, whereas no connections were observed between the GFP- and mCherry-expressing vascular-like networks (Fig. 4b). Throughout the continued culture of the SH organoid on the chip, the EGFP- and mCherry-expressing vascular-like networks were connected as vascular-like networks by day 12 (Fig. 4c). 3D construction using z-stack images of the SH organoid-on-a-chip on day 12 showed vascular-like network connections between GFP- and mCherry-expressing HUVECs (Fig. 4d). For detailed observations, we employed confocal microscopy. A 3D image created from z-stack data showed that the EGFP-expressing vascular-like networks overlapped with the mCherry-expressing vascular-like networks on day 12 (Fig. 4e). To evaluate vascular connectivity, we analyzed the xy, xz, and yz planes and created an ortho image. We found co-localization of GFP and mCherry signals, suggesting possible connections between HUVECs grown from peripheral channels and the SH organoid-derived HUVECs. These findings demonstrated that the SH organoids maintained vascular-like networks more effectively than the sclerotome spheroids on the chip. Given that hESC-derived sclerotomes have differentiation potentials into skeletal cell types, a series of our data suggest that the SH organoid-on-a-chip method can, at least partially, recapitulate vascular interactions that occur at the initial stage of endochondral ossification.

**Fig. 4.**
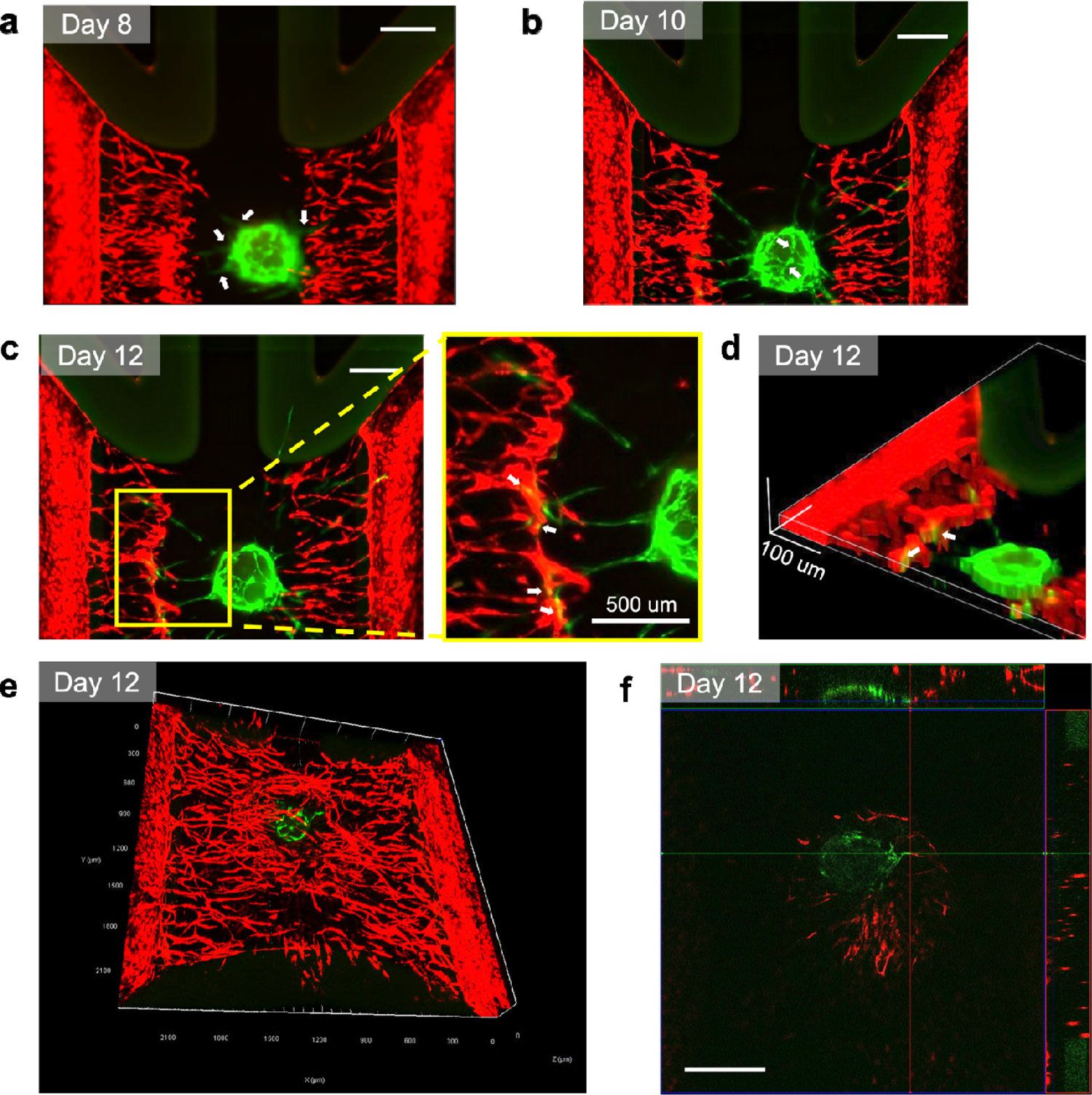
Interaction between SH organoid and peripheral vascular-like network, represented by the connection between mCherry-expressing HUVECs grown from peripheral channels and the SH organoid-derived EGFP-expressing HUVECs. (a) Fluorescent microscopy images showing vascular-like networks on day 8 (a), day 10 (b) and day 12 (c). White arrows show the vascular-like networks. In (c), yellow box is zoomed up on the right side. (d) 3D reconstruction of the vascular-like network images using FIJI. (e) 3D confocal image of vascular-like networks on day 12. (f) xyz-view of SH organoid-on-a-chip captured with confocal microscopy. White arrows show the vascular-like network connections. For (a), (b), (c), and (f) scale bar: 500 μm.

### 3.3 Validation of endochondral ossification in the SH organoid-on-a-chip

To examine whether the SH organoid-on-a-chip underwent the process of endochondral ossification, we performed histological analysis of the SH organoids cultured for 7 days on the chip, following 14-day culture in 96-well plates (day 14+7 SH organoid-on-a-chip). H&E staining highlighted the SH organoid and its surrounding gel matrix in the day 14+7 SH organoid-on-a-chip (Fig. 5; the organoid is indicated by the yellow dashed line). Immunohistochemistry revealed weak expression of SOX9 in both the central and outer layers of the day 14 SH organoids (without additional on-chip culture), with no COL1 expression detected. In the day 14+7 SH organoid-on-a-chip, SOX9 was highly expressed in the center of the SH organoid, while COL1 was expressed in the outer layer of the organoid (Fig. 5). These results suggest that the SH organoid-on-a-chip can recapitulate the initial stage of endochondral ossification, in which SOX9-positive mesenchymal condensations are surrounded by the COL1-positive fibrous layer.

**Fig. 5.**
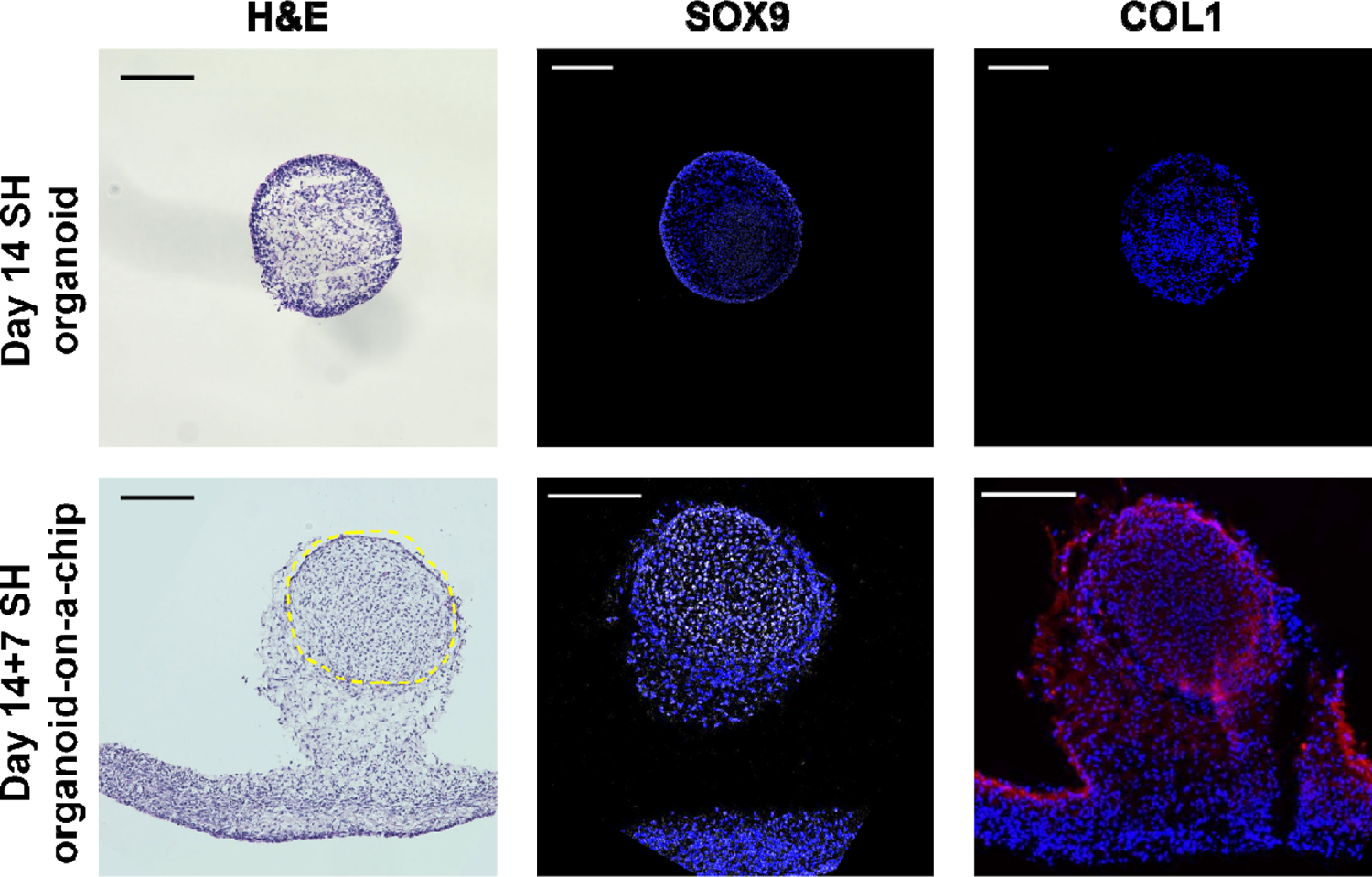
Histological analysis of SH organoid-on-a-chip. The SH organoids were initially cultured in 96-well plates for 14 days (Day 14 SH organoid) before being further cultured on a chip for an additional 7 days (Day 14 + 7 SH organoid-on-a-chip). In the H&E staining, the yellow dashed area indicates the location of the SH organoid, while the remaining tissue is fibrin gel. For SOX9 and COL1 immunostaining, SOX9 is visualized in white, COL1 in red, and nuclei are stained with DAPI, appearing blue. Scale bar: 200 μm.

## 4. Discussion

This study had 3 major findings. First, the fibrin gel formed better vascular-like networks than the collagen-I gel on the chip. Second, the SH organoid can form a vascular-like network in the organoid and interact with the peripheral vascular-like networks. Third, the SH organoid-on-a-chip maintained the condition of the vascular-like networks better than the sclerotome spheroid did. Lastly, the SH organoid-on-a-chip mimicked the initial stages of endochondral ossification. Collectively, we developed the SH organoid-on-a-chip system, which incorporates a 3D vascular-like network to model the initial stages of endochondral ossification.

The ECM plays a crucial role in vascular-like networks, and fibrin gel is better than collagen-I gel in our system. Although previous studies have used collagen-I gel for hepatic spheroids in the same organ-on-a-chip system (Organoplate^**®**^ Graft) [18], we observed shrinkage of the collagen-I gel during the induction of vascular-like networks with an angiogenic cocktail. Consistent with our results, Jung et al. applied the same fibrin gel to the same chip system with different contents and formed a vascular-like networks using primary human pulmonary microvascular endothelial cells, fibroblasts, and pericytes [17]. Nashimoto et al. used fibrin–collagen gel in a custom-made chip system [19]. However, maintaining the vascular-like networks on a chip for long term remains challenging. As shown by Bonanini, et al., [18] we observed vascular degradation during culture. Our observations suggest that after the grafting of organoids, cells from the organoid start migrating into the matrix and quickly cover the ECM during culture. This has a negative impact on the vascular-like network, although this phenomenon was uncontrollable in this system [20]. Therefore, to maintain vascular-like networks, it is important to inhibit the migration of cells from organoids. Additional studies are needed to develop a better system or ECM to prevent cell migration from the organoid tissue.

In this study, we developed the SH organoid: mixing hESC-derived sclerotome and HUVECs. This method helps the formation of vascular-like networks within the organoid. In addition, in the SH organoid on a chip system, vascular-like networks were observed between vascular-like networks from the SH organoid and that from the surrounding area in a 3D environment. This was inspired by other studies suggesting that using pre-vascularized organoids can improve the vascularization in the organoid as well as supporting vascular integration with the peripheral vascular-like networks on a chip. In addition, the use of a pre-vascularized organoid can increase the possibility of on-a-chip anastomosis [18–19–21]. We successfully created a similar vascular-like network using SH organoid-on-a-chip system.

The histological analysis suggested that our system recapitulated certain aspects of endochondral ossification. The specific expressions of SOX9 and COL1 in the SH organoid-on-a-chip are consistent with the initial stage of endochondral ossification[22], suggesting that this model allows the investigation of mechanisms underlying vascular interactions with mesenchymal condensation and the subsequent early chondrogenesis. However, to achieve modeling of the whole process of endochondral ossification, the following 3 limitations need to be overcome. First, vascular invasion into cartilage tissue is necessary for endochondral ossification. In this study, we used hESCs-derived sclerotome that includes skeletal progenitors; however, those cells have not been differentiated into mature skeletal cells yet. Given that hypertrophic chondrocytes secrete angiogenic factors for vascular interactions, maturation of the SH organoid is the next challenge. Second, the perfusion of vascular-like networks in organoids must be developed. Although we confirmed the perfusion of the vascular-like networks formed by peripheral channels, perfusion into the organoid has not yet been achieved. One possible solution may be to develop a chip device that ensures close contact of organoids with surrounding vascular-like networks, as demonstrated by Quintard et al. [21]. Additionally, as confirmed by Nashimoto et al., increasing the size of the pores in the peripheral channels can induce wider vascular-like networks (>100 μm diameter), which potentially enhance the interaction and eventually invade the organoid-on-a-chip [19]. Finally, although our study demonstrates the successful formation of a vascular-like network on a chip, there remains a gap in the long-term culture of SH organoids on a chip. As previous studies using the same chip system showed a limitation in culture periods for different cell types [18–20], we believe that the improvement or development of a chip system is required. Recently, several circulatory systems have been developed for long-term organoid culture [21]. Thus, these systems are promising for long-term culture applications.

## 5. Conclusions

This study partially provides a proof-of-concept for modeling the vascular interaction that occurs at the initial stage of human endochondral ossification. Although further improvements are required, the modeling system may contribute to a better understanding of human bone development and diseases, potentially paving the way for the discovery of novel therapeutic targets in the future.

## Abbreviations

HUVECs: human umbilical vein endothelial cells
Scl: Sclerotome
SH: sclerotome and HUVECs
VEGF: vascular endothelial growth factor
ECM: extracellular matrix

## Acknowledgments

We thank Sakurako Hayashi, Dahlia Eldeeb, Shant Nepal, Priscillia Christiany, Ireen James, Jinyan Si, Maki Kurihara, Dr. Shinse Fujita, Chie Kataoka, Asuka Uchida, and Nozomi Nagumo at The University of Tokyo for providing technical assistance. This work was supported by Grants-in-Aid for Scientific Research from the Japan Society for the Promotion of Science (JSPS: 21H04952), Rising Star Award from American Society for Bone and Mineral Research, the Japan Science and Technology Agency (JST) FOREST Program (JPMJFR225N) and JST ERATO program (JPMJER2401). This work utilized the core research facility of the Center for Disease Biology and Integrative Medicine at The University of Tokyo Graduate School of Medicine. A.K. was supported by the Japan International Cooperation Agency (JICA) Friendship programs.

## Supplementary data

**Fig. S1.**
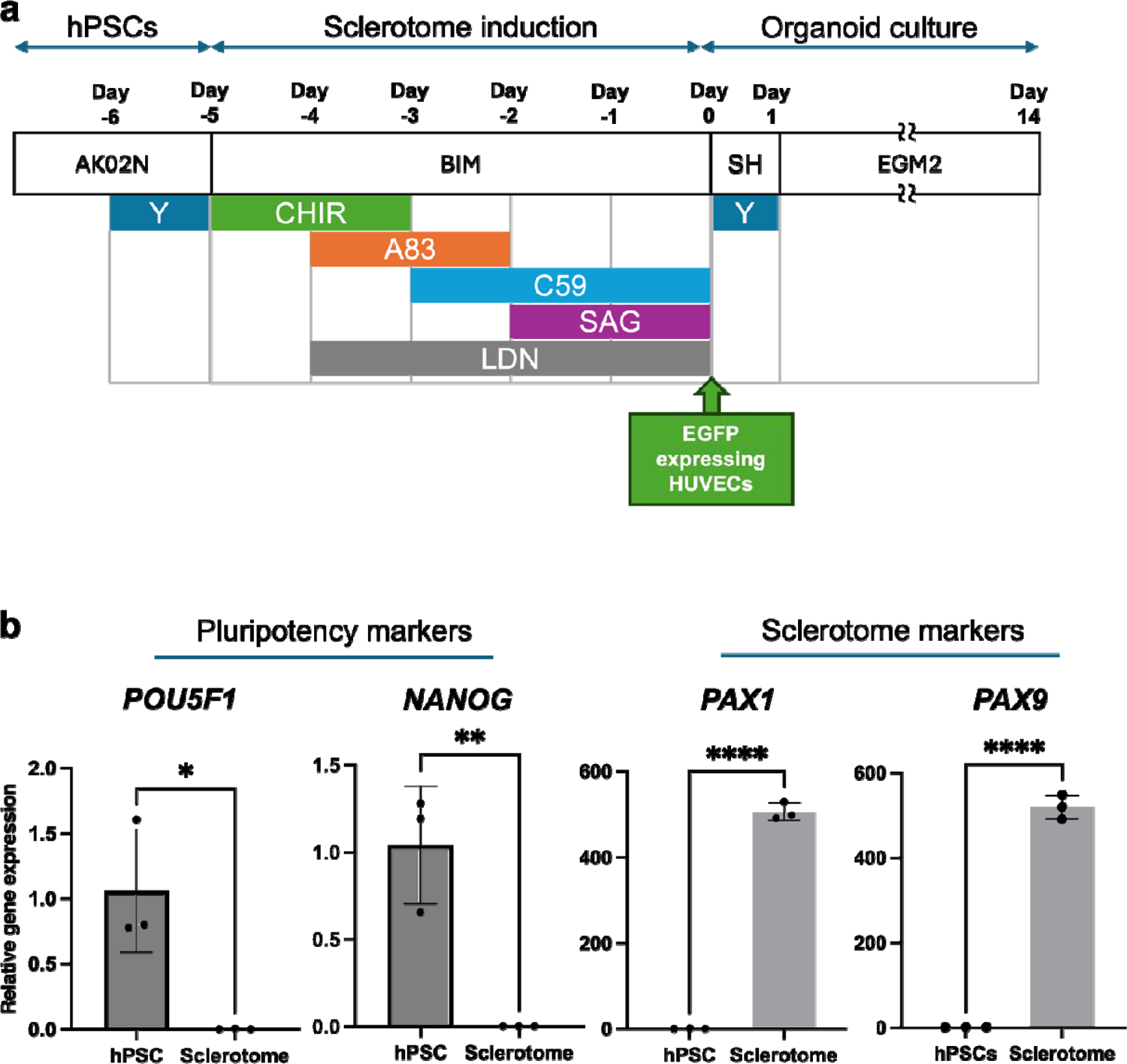
Experimental scheme of the sclerotome induction and organoid culture. (a) Sclerotome spheroids are induced form hESCs-derived sclerotome cells using different small molecules, but SH organoid is the mixture of sclerotome and EGFP expressing HUVECs. (b) Relative gene expression of pluripotent markers on day -6 and sclerotome markers on day 0, determined by RT-qPCR. Student’s t-test was used to analyze the data, “*” indicates statistical significance (*p < 0.05; **p < 0.01; ****p < 0.0001).

**Fig. S2.**
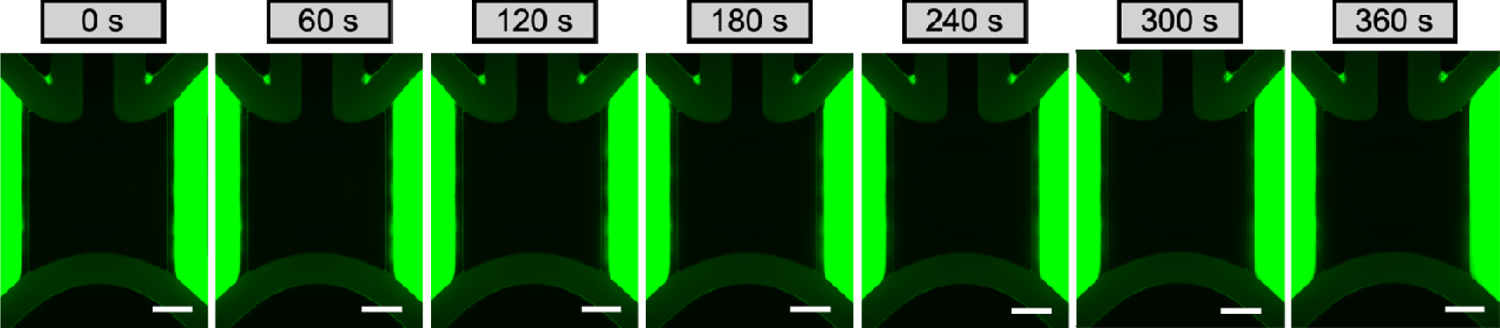
Confirmation of no perfusion of 70 kDa FITC-dextran in the absence of vascular-like networks. Scale bar: 500 μm.

